# Mutations in MreB suppress β-lactam sensitivity upon c-di-AMP accumulation in *Listeria monocytogenes*

**DOI:** 10.64898/2026.05.14.724990

**Authors:** Shreya Kumar, Hung Dang, TuAnh N. Huynh

## Abstract

Cyclic di-AMP (c-di-AMP) is an essential second messenger in *Listeria monocytogenes*, but its accumulation is detrimental as it disrupts cell wall homeostasis and attenuates virulence. The mechanisms underlying this toxicity remain poorly understood. To understand the molecular basis of this toxicity, we performed a forward genetic screen to identify suppressor mutations that restore β-lactam resistance in a Δ*pdeA* Δ*pgpH* (Δ*PDE*) mutant, which accumulates high c-di-AMP and is susceptible to cell wall-targeting β-lactam antibiotics. We found that the majority of suppressor mutants carried mutations in the *mreB* gene, which encodes the bacterial actin-like cytoskeletal protein, MreB, that directs lateral peptidoglycan synthesis during cell elongation. These mutations restored β-lactam resistance and ex vivo virulence while still retaining high intracellular c-di-AMP levels. Microscopy analyses indicate that these suppressor mutations reduce MreB activity, as evidenced by cell widening, and that they phenocopy sublethal treatment with the MreB inhibitor A22. Consistently, A22 treatment also rescued β-lactam sensitivity in the Δ*PDE* mutant, supporting a functional link between MreB activity and c-di-AMP toxicity. Mechanistically, c-di-AMP accumulation impaired cell division/septation and reduced peptidoglycan synthesis under cell wall stress, whereas MreB mutations restored both transglycosylation and transpeptidation activities and promoted cell division. These effects were independent of potassium homeostasis, suggesting a distinct pathway linking c-di-AMP to cell wall regulation in *L. monocytogenes*. Together, our findings demonstrate that dysregulated MreB activity contributes to cell wall defects at elevated c-di-AMP levels and highlight the importance of coordinating cytoskeletal dynamics with cell division to maintain cell envelope integrity.

## INTRODUCTION

The Gram-positive bacterial pathogen *Listeria monocytogenes* requires c-di-AMP for growth in rich laboratory media and during infection. *L. monocytogenes* synthesizes c-di-AMP by a single diadenylate cyclase, DacA, and hydrolyzes it by the phosphodiesterases PdeA (also called GdpP in other bacteria) and PgpH (1, 2). The Δ*pdeA* Δ*pgpH* mutant (hereafter denoted as Δ*PDE*), which accumulates c-di-AMP while exhibiting largely normal growth in rich broth media, is significantly attenuated for virulence (3). The essentiality of c-di-AMP in *L. monocytogenes* is attributed to its role in regulating the stringent response, since the Δ*dacA* mutant accumulates ppGpp, which inhibits growth (4). By contrast, the mechanisms for c-di-AMP toxicity in *L. monocytogenes* are not fully defined.

In some Gram-positive Firmicutes, mutants with elevated c-di-AMP levels show increased resistance to cell wall-targeting antibiotics, such as β-lactams (5–8). Indeed, loss-of-function mutations in GdpP are strongly associated with β-lactam resistance in *Enterococcus faecalis* and in clinical isolates of *Staphylococcus aureus* (9, 10). These cell wall phenotypes have been attributed to reduced cell turgor, due to inhibition of potassium and osmolyte transport by cdA. Curiously, the *L. monocytogenes* Δ*PDE* exhibits a loss in cell wall integrity despite a deficiency in cytoplasmic K^+^ (11). Compared to the wild-type (WT) strain, the Δ*PDE* mutant is sensitive to cell wall-targeting antibiotics such as D-cycloserine, lysozyme, and β-lactams, and exhibits a deficiency in cytoplasmic peptidoglycan precursors (11).

Peptidoglycan (PG) is the major component of the bacterial cell wall. PG precursors are synthesized in the cytoplasm by the activities of Mur and Ddl enzymes, generating UDP-MurNAc-pentapeptide as the final soluble precursor. This precursor is then ligated with undecaprenol to generate lipid II, which is flipped across the membrane for assembly into the PG. In the extra-cytoplasmic space, transglycosylases polymerize PG precursors into alternating GlcNac and MurNac subunits, and transpeptidases intermittently crosslink pentapeptide stems from opposite polymer strands (12). Rod-shaped bacteria synthesize PG both for cell elongation and division (13, 14). Lateral PG synthesis is largely dependent on the Rod complex, comprised of penicillin binding protein 2A (PBP 2A), RodA, RodZ, MreC, MreD, and MreB. RodA polymerizes PG precursors, and PBP2A crosslinks glycan strands (12). MreB polymerizes into filaments that interact with RodZ, and spatially guide Rod activity to maintain the rod shape as the cell wall expands (12, 15, 16). Although all Rod complex components are required to maintain the rod shape, only MreB is unique to rod-shaped bacteria and is absent in coccoid bacteria. MreB is essential for optimal growth and cell morphology in rod-shaped bacteria (14–16). However, hyperactive MreB is also toxic, as *mreB* overexpression diminishes growth and causes cell lysis (17–20). Under cell wall stress and PG precursor depletion, MreB depolymerizes and dissociates from the membrane (21).

Here, we employed forward genetics to define the molecular basis of cell wall defects associated with c-di-AMP accumulation in *L. monocytogenes*. We chemically mutagenized the Δ*PDE* strain and screened for mutants that were restored for β-lactam resistance while retaining high c-di-AMP levels. Our genetic screen repeatedly identifies mutations in MreB that restore both β-lactam resistance and ex vivo infection by the Δ*PDE* strain, without correcting c-di-AMP levels. These mutants exhibited widened cells, consistent with a reduced MreB activity. Sublethal β-lactam treatment inhibited cell division in *L. monocytogenes*, with a visibly pronounced effect in the Δ*PDE* strain. At high c-di-AMP levels, MreB mutations rescued septum formation, and overexpression of *pbp B2*, the main penicillin-binding protein involved in cell division, modestly increased β-lactam resistance. Importantly, at normal c-di-AMP levels, MreB mutations did not increase β-lactam resistance but did cause aberrant cell morphology. Together, our study suggests that c-di-AMP accumulation inhibits cell division in *L. monocytogenes* and that cell wall defects are a major contributor to virulence attenuation in the Δ*PDE* strain.

## RESULTS

### Genetic screen identifies MreB mutations that rescue β-lactam resistance in *L. monocytogenes* Δ*PDE*

In several bacteria, c-di-AMP accumulation increases resistance or tolerance to cell wall-targeting antibiotics, such as β-lactams. By contrast, the *L. monocytogenes* Δ*PDE* mutant is β-lactam-sensitive (11). To identify the molecular basis of the cell wall defects in Δ*PDE*, we performed a forward genetic screen to identify suppressor mutations that rescue its β-lactam sensitivity. Although we could easily obtain spontaneous suppressor mutations by passaging Δ*PDE* under stress, the vast majority were in the c-di-AMP synthase DacA, reflecting a high selective pressure for c-di-AMP homeostasis (22–24). As an alternative approach, we performed chemical mutagenesis with EMS (ethyl methanesulfonate), which induces GC → AT transitions in DNA (25, 26). Following treatment of the Δ*PDE* strain with EMS, we screened for cefuroxime-resistant colonies on BHI agar containing cefuroxime at the minimum inhibitory concentration for the parent Δ*PDE* strain. Mutant colonies, abbreviated as Cef^R^, arose at the frequencies of ∼ 10^-4^. Upon subsequent growth in BHI broth containing cefuroxime, approximately 20% of Cef^R^ candidates were confirmed to exhibit WT-level resistance. We selected 40 mutants for LC-MS/MS quantification of c-di-AMP and found that 70% exhibited reduced c-di-AMP levels compared to the Δ*PDE* strain. Among the remaining mutants that retain high c-di-AMP levels, we selected 10 for whole-genome sequencing and identified 1-3 mutations in each. Intriguingly, 8 out of 10 Cef^R^ suppressor mutants harbored mutations in MreB, with three of them having only a single mutation in MreB (**Figs. 1A-B and 2**).

**Figure 1:**
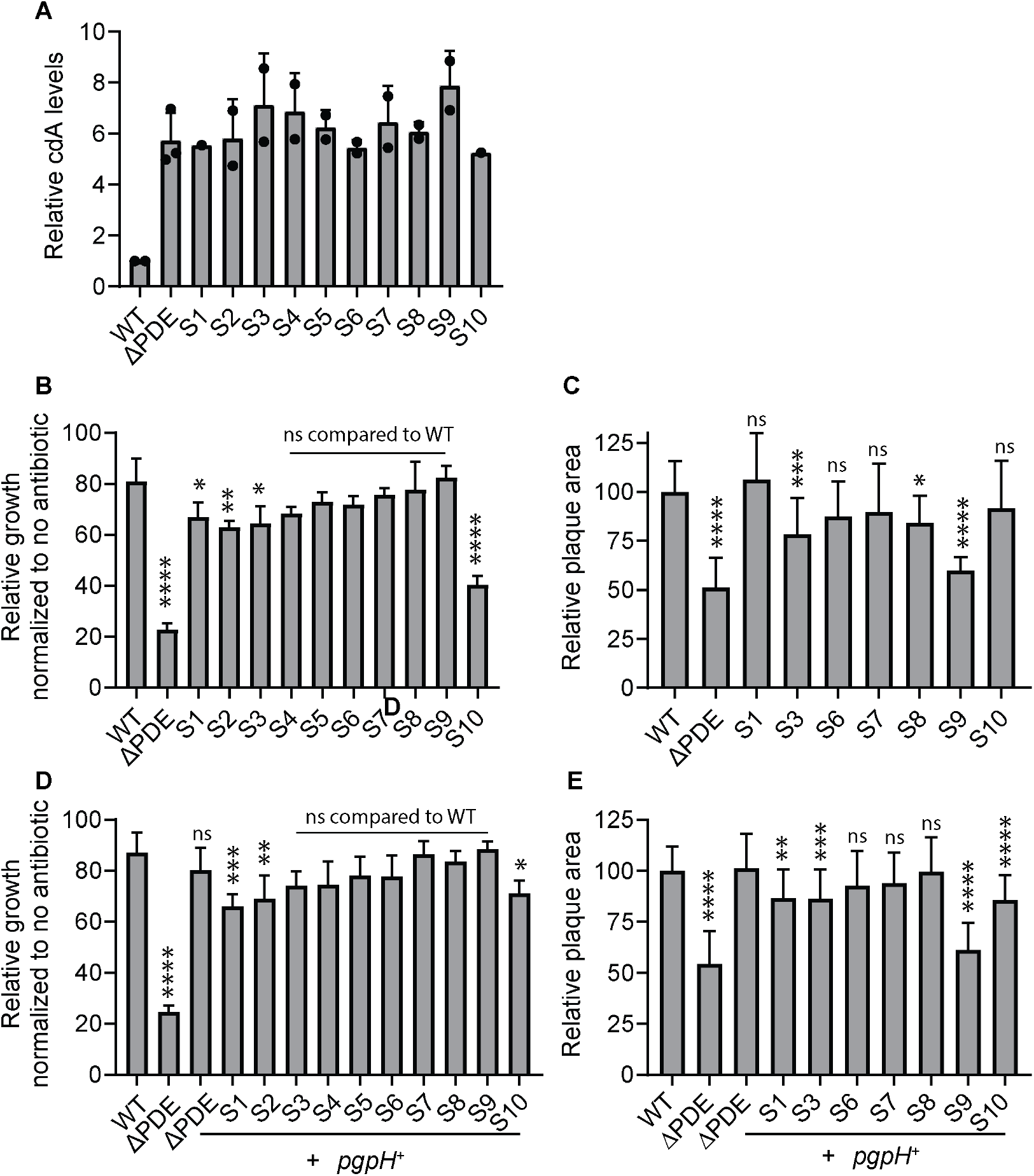
Suppressor mutants are restored for β-lactam resistance and ex vivo infection in the Δ*PDE* genetic background. **A**. Quantification of c-di-AMP levels by LC-MS/MS, with C^13^ N^15^-c-di-AMP as an internal standard, and normalized to bacterial biomass (OD_600_ X Volume). **B and D**. Relative growth rates of *L. monocytogenes* in BHI + 0.625 µg/mL cefuroxime, normalized to growth rates in BHI only for each strain. **C and E**. Plaque formation at 4 days post-infection of L2 fibroblasts. Plaque sizes were quantified by ImageJ. In each experiment, the plaque size of each plaque by each strain was normalized to the average WT plaque size. Error bars show standard deviations. Statistical analyses were performed by one-way ANOVA, with multiple comparisons for the indicated pairs: ns, non-significant; *, P < 0.05; **, P < 0.01; ***, P < 0.001; ****, P < 0.0001

In addition to β-lactams, the Δ*PDE* strain is also sensitive to other cell wall-targeting antibiotics, including D-cycloserine, bacitracin, and lysozyme (11) (**Fig. S1**). Almost all suppressor mutants remained sensitive to these antibiotics. As an exception, mutant S10 was restored for lysozyme resistance. A major mechanism of lysozyme resistance in *L. monocytogenes* involves the deacetylation and O-acetylation of peptidoglycan. In *Bacillus subtilis*, WalR inhibits the transcription of the deacetylase Pgd (27, 28). While the WalR regulon in *L. monocytogenes* is not yet defined, it is conceivable that a *walR* mutation in S10 derepresses *pgd*, leading to increased PG deacetylation and, consequently, increased lysozyme resistance.

Finally, the Δ*PDE* strain also shows reduced virulence in a mouse model of intravenous infection. This defect can be reported in an ex vivo plaque formation assay upon infection of L2 fibroblast cells. Strikingly, nearly all Cef^R^ suppressor mutants were also restored for plaque formation, except mutant S9 (**Fig. 1C**).

### Suppressor effect is specific to high c-di-AMP levels in *L. monocytogenes*

Nine out of ten suppressor mutants (S1 – S9) were rescued for cefuroxime resistance almost to the WT level (**Fig. 1B**). Mutants S1 – S8 all harbor MreB mutations, and mutant S9 harbors a mutation in RpoD, an RNA polymerase subunit. Mutant S10, which harbors a mutation in the response regulator WalR, exhibited an intermediate resistance between WT and Δ*PDE* (**Figs. 1B and 2**). To test the effects of these mutations at normal c-di-AMP levels, we complemented S1 – S10 with PgpH, a major c-di-AMP phosphodiesterase in *L. monocytogenes*. These complemented strains were not more cefuroxime resistant than WT (**Fig. 1D**). Indeed, mutants S1, S2, and S10 remained sensitive upon complementation with PgpH. Similarly, plaque formation by these suppressor mutants was unaffected by PgpH complementation (**Fig. 1E**). Therefore, the suppressor effect was specific to high c-di-AMP levels.

### A reduced MreB activity increases β-lactam resistance at high c-di-AMP levels

Given the repeated isolation of MreB mutations in our genetic screen, we next sought to identify how they restore β-lactam resistance in the Δ*PDE* strain. We specifically focused on mutants S3, S6, and S8, since each harbored a single MreB mutation in its genome (**Fig. 2**). Based on structural homology of *L. monocytogenes* MreB with the crystal structures of *Thermotoga maritima* MreB (PDB 1JCG), mutant S3 harbors an A46V mutation in the protomer polymerization surface, mutant S6 harbors a G108E mutation within the ATP-binding pocket, and mutant S8 harbors a V47I mutation near the interaction surface with RodZ (29, 30) (**Fig. 2**).

**Figure 2:**
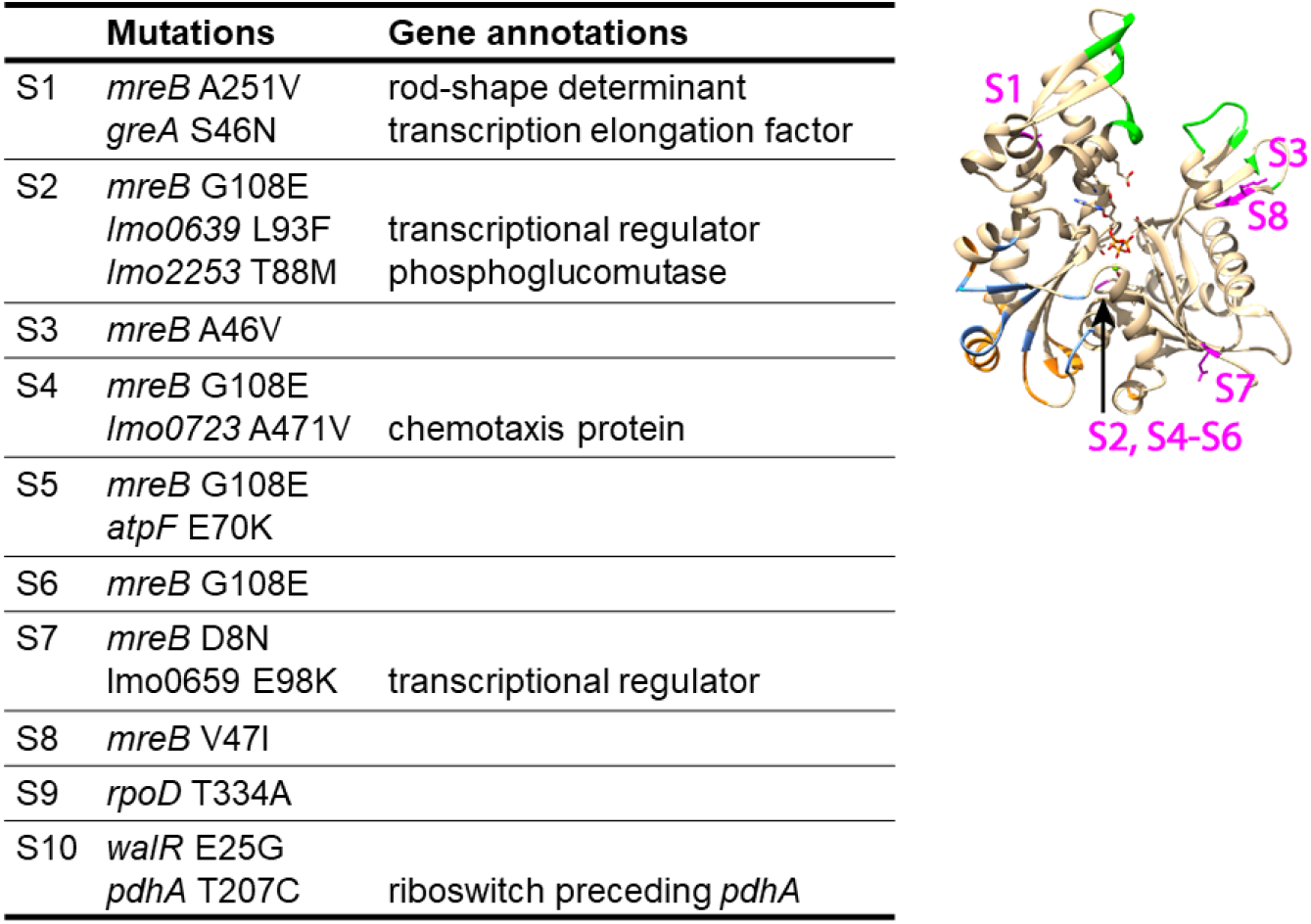
Mutations in MreB are frequently isolated among Δ*PDE* suppressor mutants. **A**. Mutations in each suppressor mutant (S1 – S10), all in the Δ*pdeA* Δ*pgpH* (Δ*PDE*) genetic background. **B**. MreB mutations mapped on the crystal structure of *Thermotoga maritima* MreB (PDB 1JCG).

MreB is an essential component of lateral peptidoglycan synthesis in rod-shaped bacteria. Loss-of-function mutations in MreB cause cell rounding, and a reduced *mreB* gene expression increases cell width (16, 31). Compared with WT and Δ*PDE*, all three suppressor mutants (S3, S6, and S8) exhibited widened cells, both under normal growth and in the presence of cefuroxime, consistent with a reduced MreB activity (**Fig. 3**). To recapitulate the effect of MreB mutations, we used compound A22, which competitively inhibits MreB enzymatic activity by binding near the ATP nucleotide binding site (29). Because A22 can be toxic to bacterial growth, independent of MreB inhibition (32), we first determined a suitable A22 concentration with minimal impacts on *L. monocytogenes* growth. At 50 µg/mL, A22 had a negligible effect on *L. monocytogenes* growth, but still widened both WT and Δ*PDE* cells, and not the S6 mutant that has an MreB G108E mutation near the ATP-binding site (**Fig. S2**). In addition, A22 treatment substantially increased cefuroxime resistance in Δ*PDE*, while sensitizing WT (**Fig. 4**). As controls, compound A22 did not affect cefuroxime resistance or cell width in MreB suppressor mutants (**Fig. S2** and **Fig. 4**). Combined, these data indicate that β-lactam sensitivity upon c-di-AMP accumulation is suppressed by reducing MreB function.

**Figure 3:**
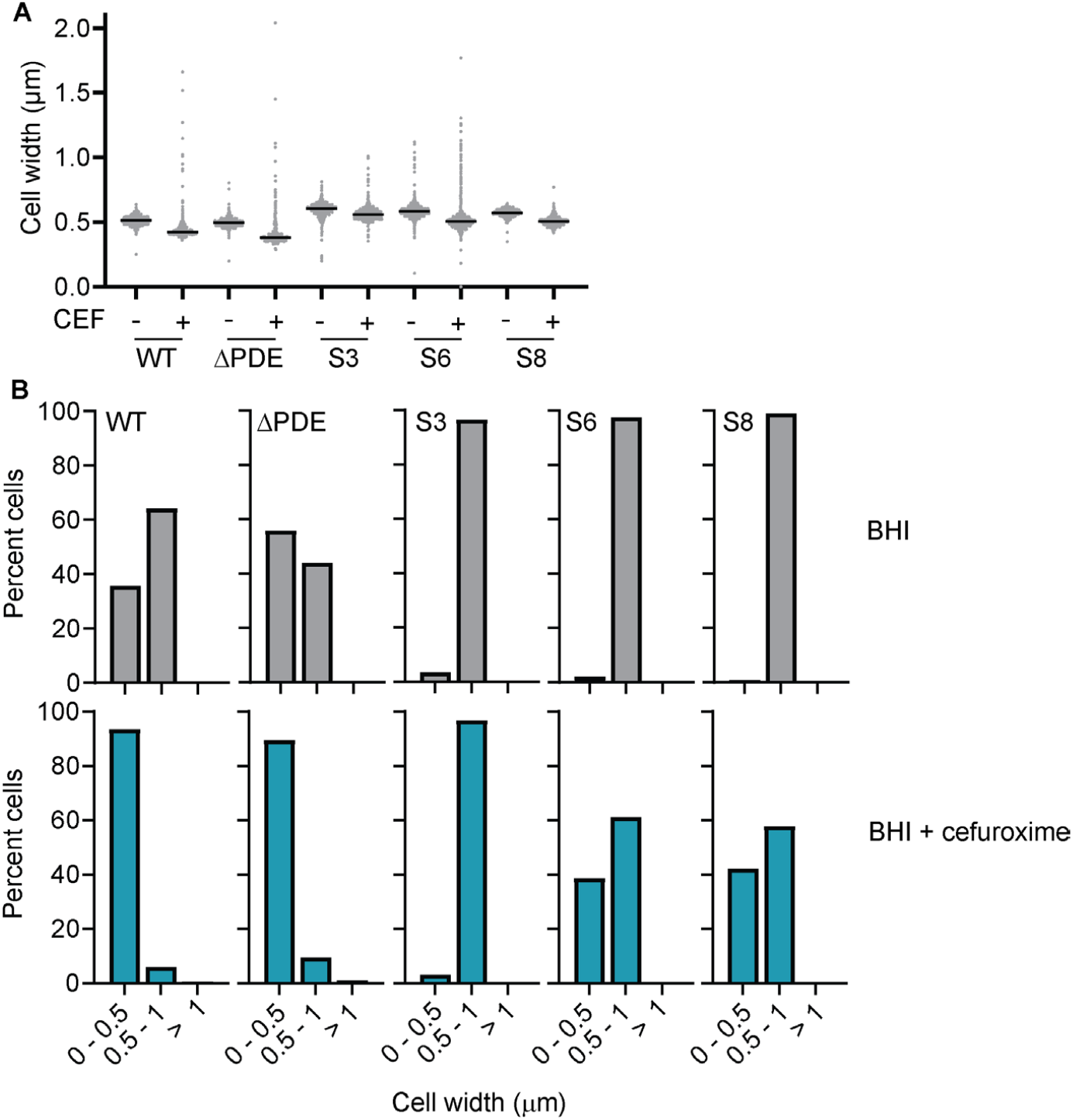
Mutations in MreB widen cells. *L. monocytogenes* strains were grown in BHI or BHI + 1.25 µg/mL cefuroxime to mid-log phase, and stained with WGA. **A**. Cell widths were measured using MorphoSegger. Between 300 and 500 cells were quantified per condition across three independent experiments. **B**. Distributions of cell width quantified in A. Width measurements were compiled into a single dataset and analyzed using a frequency distribution, with values grouped into fixed-length bins of 0.25 μm, spanning 0 to 2.25 μm. Bin boundaries were defined as left-inclusive and right-exclusive (e.g., 0.00–0.25, 0.25–0.50, etc.), and each data point was assigned to exactly one bin. Exact counts were calculated for each bin.

**Figure 4:**
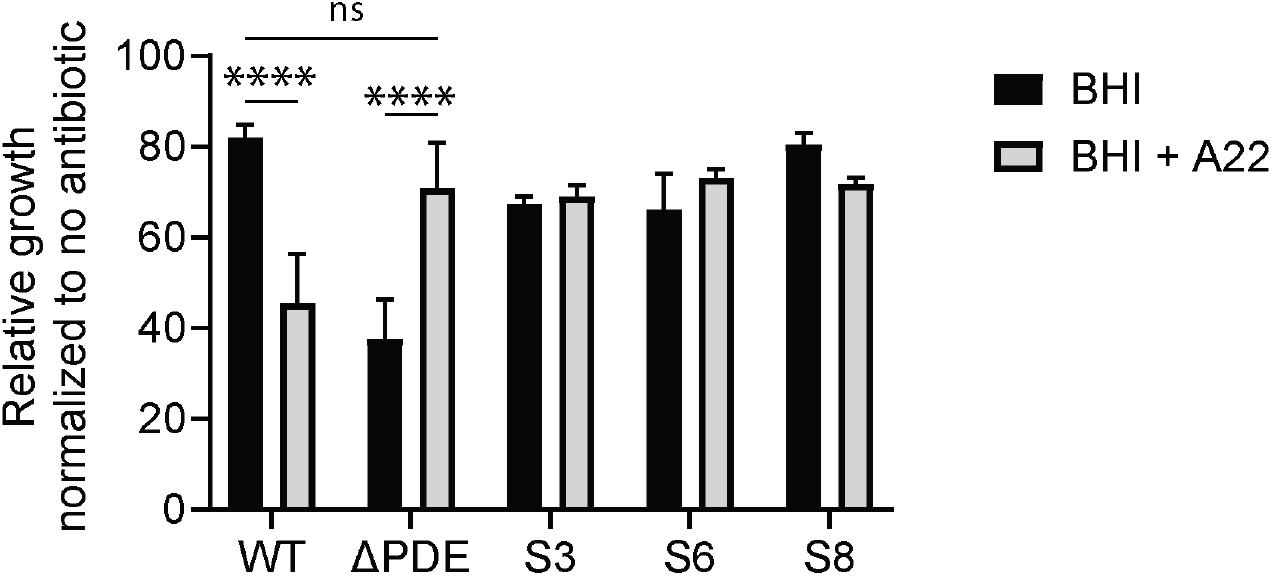
A sub-inhibitory A22 treatment increases cefuroxime resistance in Δ*PDE*. *L. monocytogenes* was grown in BHI or BHI + 50 µg/mL A22, with or without 0.625 µg/mL cefuroxime. Relative growth rates in cefuroxime were normalized to growth rates in the absence of antibiotics. Error bars show standard deviations. Statistical analyses were performed by one-way ANOVA, with multiple comparisons for the indicated pairs: ns, non-significant; *, P < 0.05; **, P < 0.01; ***, P < 0.001; ****, P < 0.0001

**Figure 4:**
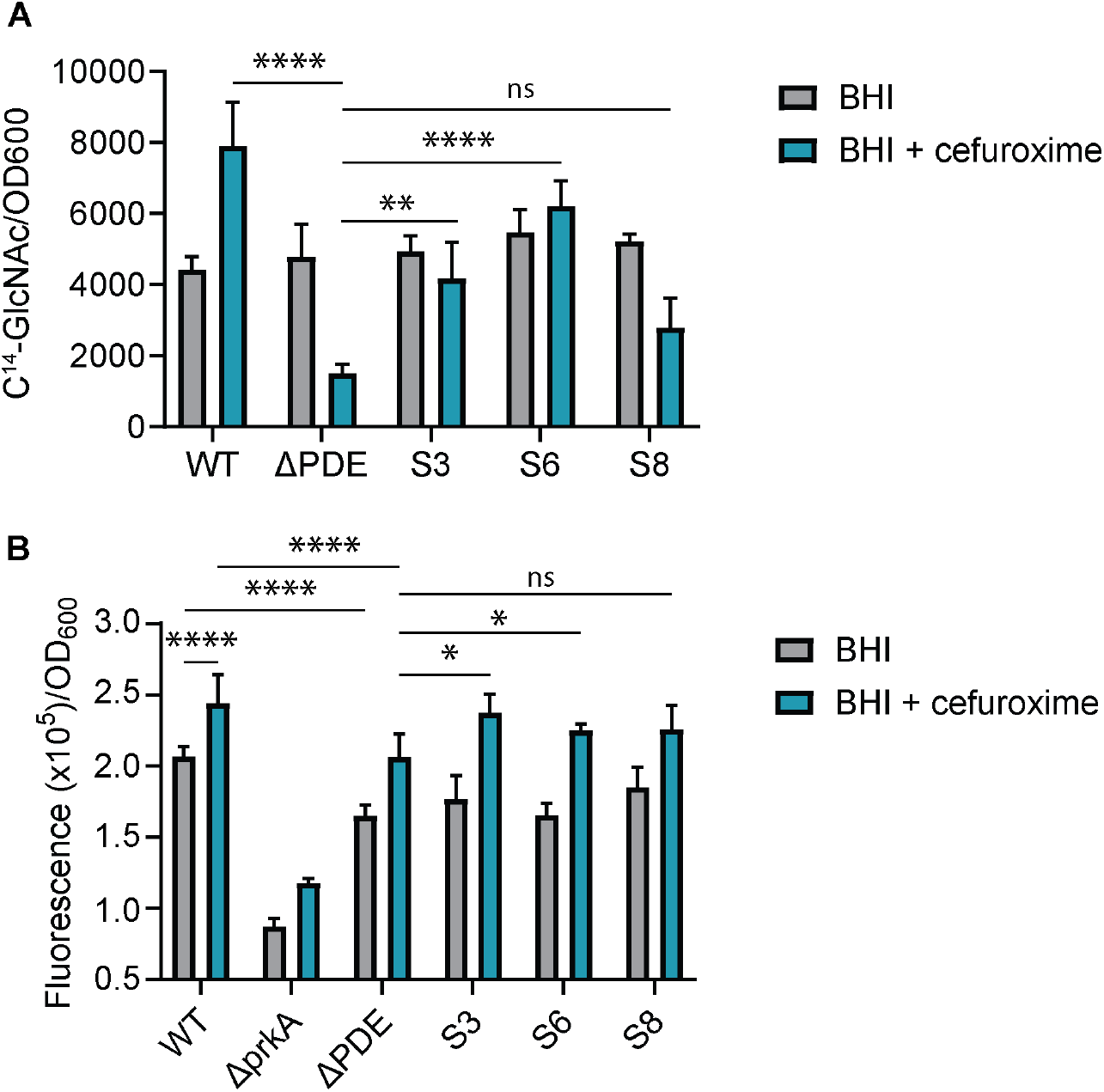
Incorporation of ^14^C-GlcNAc and EDA-DA. A. ^14^C-GlcNAc incorporation by *L. monocytogenes*. Radioactivity was normalized to OD_600_. **B**. EDA-DA incorporation by *L. monocytogenes*, detected with Alexa Fluor 488 azide. Fluorescence was normalized to OD_600_. *L. monocytogenes* cultures, for both assays, were grown in BHI or BHI + 1.25 µg/mL cefuroxime, except for Δ*prkA*, which was grown in BHI + 0.125 µg/mL cefuroxime. Error bars represent standard deviations. Data are the average of 3-5 independent experiments. Statistical analysis was performed by one-way ANOVA, with multiple comparisons for the indicated pairs: ns, non-significant; *, P < 0.05; **, P < 0.01; ***, P < 0.001; ****, P < 0.0001.

### Suppressor effect of MreB mutations is independent of K^+^ availability

A conserved function of c-di-AMP is to regulate K^+^ homeostasis, through inhibiting K^+^ import and activating K^+^ uptake. Therefore, c-di-AMP accumulation results in a net reduction in cytoplasmic K^+^ levels and cell turgor. In vitro, MreB depolymerizes at high K^+^ concentrations. In *B. subtilis*, osmotic stress induces MreB depolymerization, a process that depends on K^+^ influx. Studies in *B. subtilis* and *Streptococcus* suggested that cell wall stress induces an osmotic stress. If β-lactams similarly cause a K^+^ influx in *L. monocytogenes* to depolymerize MreB, then WT, Δ*PDE*, and MreB suppressor mutants would exhibit similar resistance at limiting K^+^ levels. However, we found that cefuroxime resistance in these strains was independent of K^+^ availability in the growth medium (**Fig. S3**).

We further quantified K^+^ levels in *L. monocytogenes* cultures grown with 0.5xMIC cefuroxime. A decrease in K^+^ was modest. Together, these data indicate that K^+^ homeostasis does not affect β-lactam sensitivity upon c-di-AMP accumulation, or explain the suppressor effect of MreB mutations.

### MreB mutations partially rescue peptidoglycan synthesis upon c-di-AMP accumulation

Peptidoglycan homeostasis is the net result of synthesis and hydrolysis, the latter of which is required to insert new materials into the expanding cell wall. For cell wall turnover, *L. monocytogenes* encodes several autolysins, some of which are induced by detergents such as Triton X-100 (33, 34). Compared to WT, the Δ*PDE* mutant exhibited increased detergent-induced autolysin activity, but this defect was not rescued in MreB suppressor mutants (**Fig. S4**).

We evaluated peptidoglycan synthesis through incorporation of ^14^C-GlcNAc, a transglycosylase reporter, and alkyne-D-alanyl:D-analine dipeptide (EDA-DA), a transpeptidase reporter (35, 36). Under normal growth, ^14^C-GlcNAc incorporation was similar among WT, Δ*PDE*, and MreB suppressor mutants. By contrast, during growth with a sub-inhibitory cefuroxime concentration (0.5xMIC of Δ*PDE*), ^14^C-GlcNAc incorporation significantly increased in WT and substantially decreased in Δ*PDE*. Suppressor mutants S3 and S6 were largely unaffected by cefuroxime treatment and, therefore, exhibited much higher ^14^C-GlcNAc incorporation than Δ*PDE* under cell wall stress (**Fig. 4A**). By quantifying EDA-DA incorporation, we found that the Δ*PDE* mutant showed reduced incorporation compared to WT under normal growth conditions, and that MreB mutations did not improve this defect. All strains increased EDA-DA incorporation under cefuroxime treatment, but Δ*PDE* activity remained low compared to WT, and this defect was substantially rescued by MreB mutations (**Fig. 4B**). As controls, the Δ*prkA* strain exhibited a severe EDA-DA incorporation defect under both conditions, as previously reported (37). Overall, these results demonstrate that MreB mutations rescued Δ*PDE* for both transglycosylase and transpeptidase reporters under cell wall stress.

### Cefuroxime inhibits *L. monocytogenes* cell division, and MreB mutations rescue division septa formation

*L. monocytogenes* is β-lactam-tolerant. During growth with a sublethal cefuroxime concentration, WT cells became thin and markedly elongated (**Figs. 5-6**). The Δ*PDE* cells also elongated but were particularly defective in septum formation, with ∼75% of cells lacking septa (**Figs. 5-7**). By contrast, suppressor mutants S3, S6, and S8, each harboring a single MreB mutation, largely maintained normal cell length and width under cefuroxime treatment, and ∼75% of their cells formed at least one division septum. We noticed that these mutants had ∼10 – 20% of cells with multiple division septa, suggesting that MreB mutations promote cell division at high c-di-AMP levels (**Figs. 5-7**). Remarkably, upon complementation with PgpH to restore c-di-AMP levels, those MreB mutants exhibited severe morphological defects, further indicating that MreB mutations specifically benefit the Δ*PDE* strain (**Fig. 5**).

**Figure 5:**
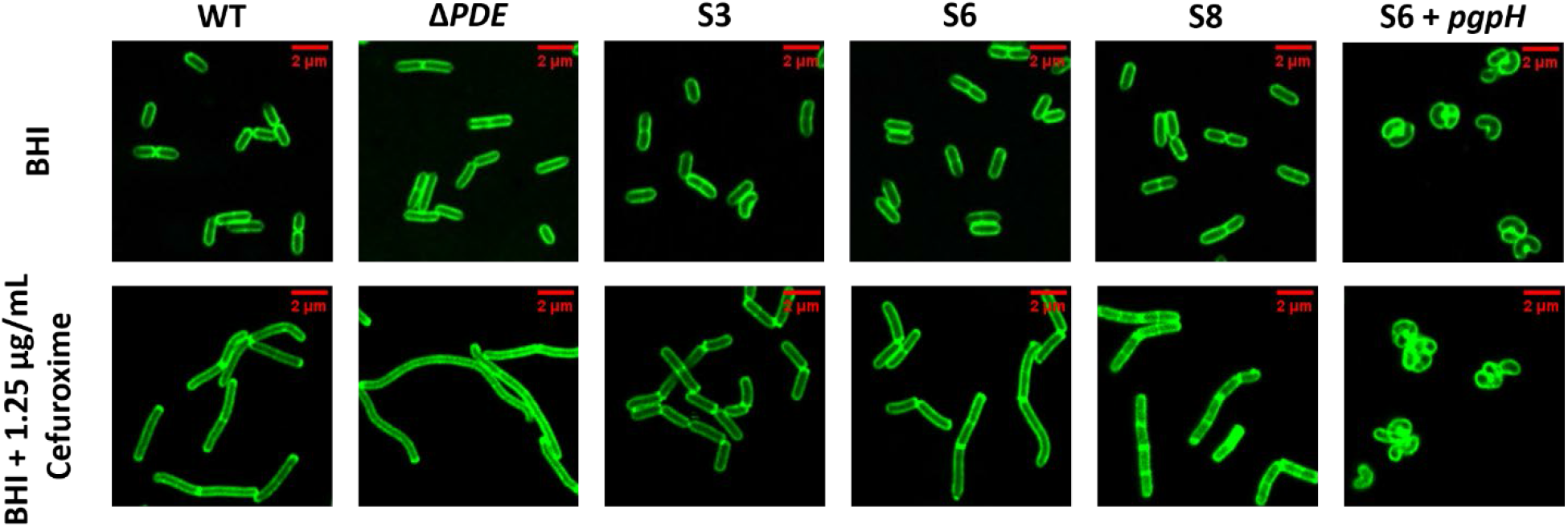
Effect of MreB Mutations on Cell morphology. *L. monocytogenes* strains were grown in either BHI or BHI with 1.25 µg/mL of cefuroxime to mid-log and stained with WGA. Scale bar = 2 µm.

**Figure 6:**
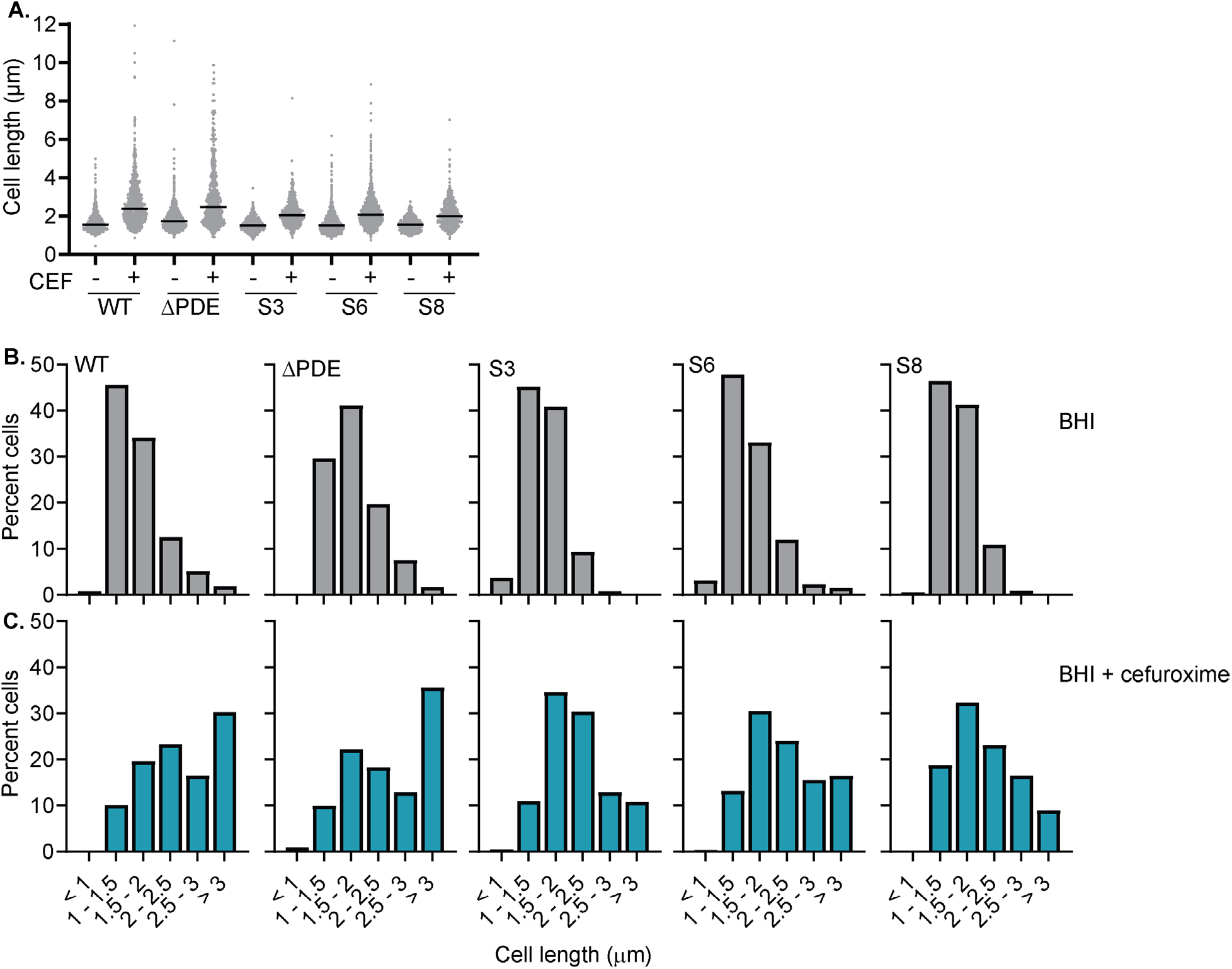
Cefuroxime treatment elongates *L. monocytogenes* cells. **A**. *L. monocytogenes* was grown in BHI or BHI + 1.25 µg/mL cefuroxime to mid-log phase and stained with WGA. Cell length was quantified with MorphoSegger. Between 300 and 500 cells were quantified per condition across three independent experiments. **B-C**. Distributions of cell length quantified in A. Length measurements were compiled into a single dataset and analyzed using a frequency distribution, with values grouped into fixed-length bins of 0.5 μm, spanning 0 to 10 μm. Bin boundaries were defined as left-inclusive and right-exclusive (e.g., 0.00–0.5, 0.5–1.0, etc.), and each data point was assigned to exactly one bin. Exact counts were calculated for each bin.

**Figure 7:**
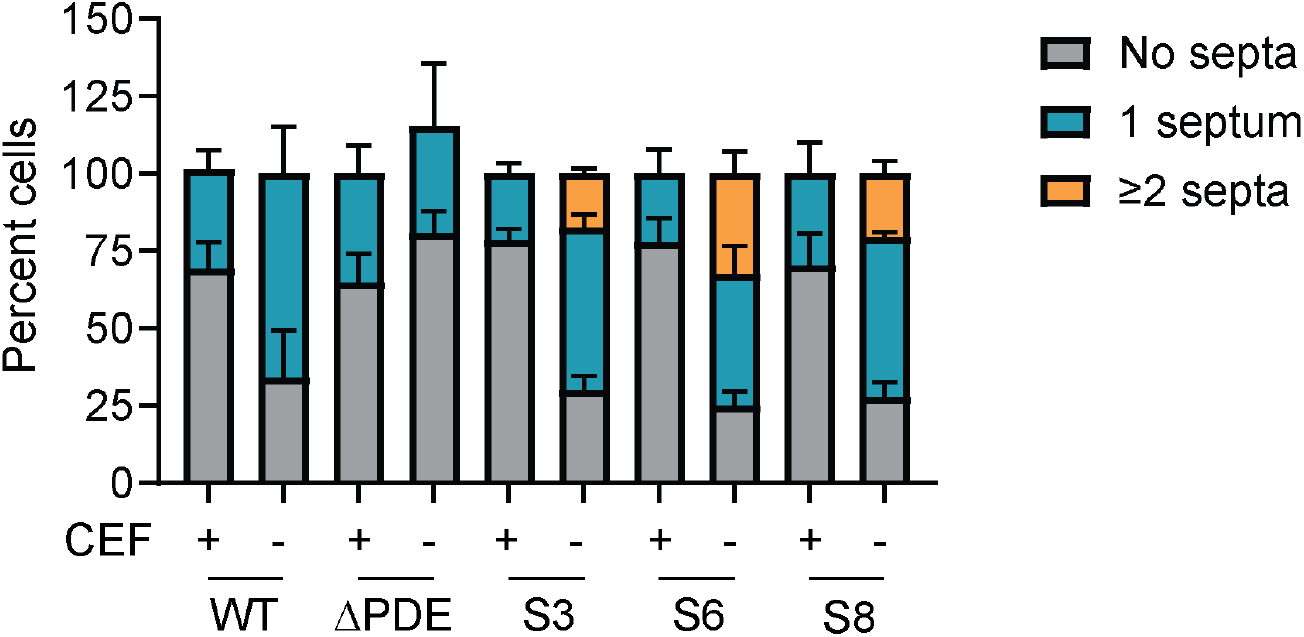
Cefuroxime treatment inhibits cell division in *L. monocytogenes*. Cells grown in BHI or BHI + 1.25 µg/mL cefuroxime were quantified for division septa in each cell. Error bars represent standard deviation.

*L. monocytogenes* encodes five penicillin-binding proteins, with the transpeptidase PBP B2 (Lmo2039) specialized for cell division (38). Genetic depletion of *lmo2039* results in elongated cells lacking division septa, reminiscent of Δ*PDE* cells under cefuroxime treatment (38). In the Δ*PDE* strain, PBP B2 over-expression significantly increased septa formation and cefuroxime resistance, although it remained more sensitive than WT (**Fig. 8**). These data suggest that PBP B2 activity is impaired at high c-di-AMP levels.

**Figure 8:**
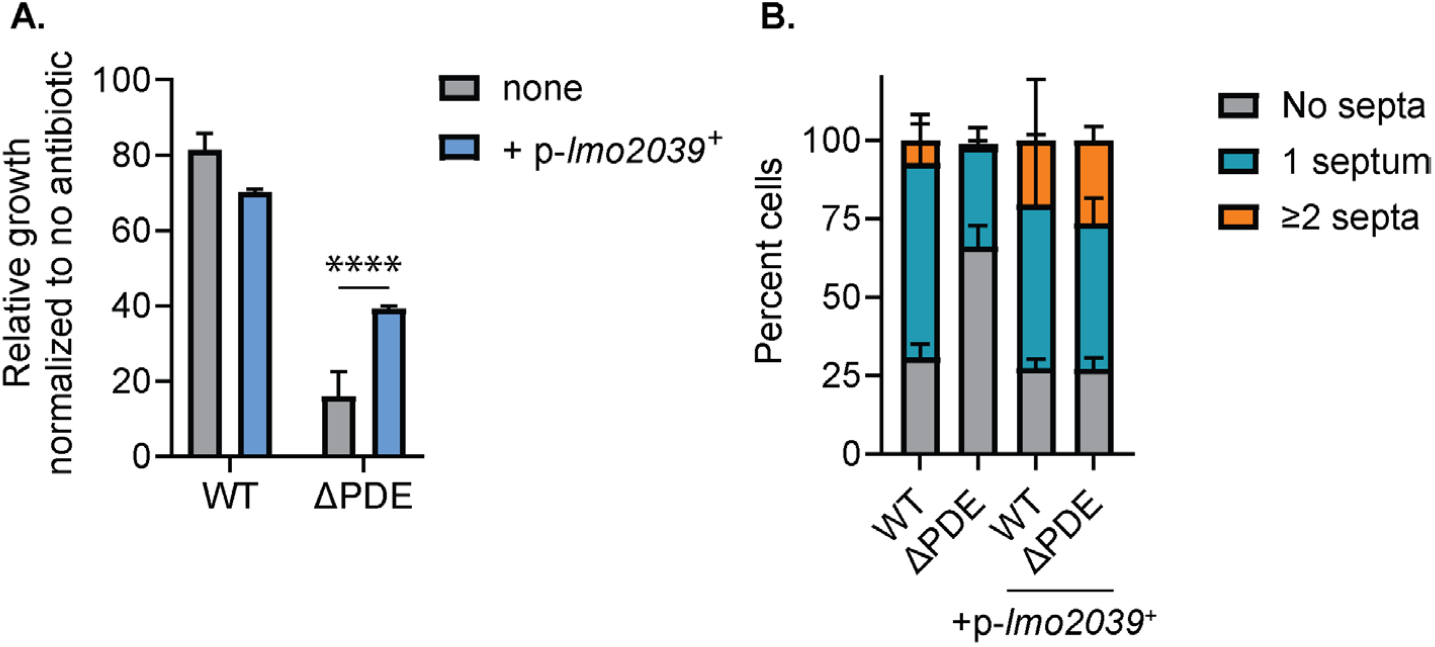
Over-expression of Pbp B2 (Lmo20239) partially rescues cefuroxime resistance and division septum formation in the Δ*PDE* strain under cell wall stress. **A**. *L. monocytogenes* was grown in LSM + 6 µg/mL cefuroxime, and growth was normalized to that in LSM only for each strain. **B**. Cells grown in BHI or BHI + 1.25 µg/mL cefuroxime were quantified for division septa in each cell. Data are the average of 5-6 independent experiments. Error bars represent standard deviation. Statistical analysis was performed by one-way ANOVA, with multiple comparisons for the indicated pairs: ns, non-significant; *, P < 0.05; **, P < 0.01; ***, P < 0.001; ****, P < 0.0001.

## DISCUSSION

C-di-AMP has been known for decades to regulate bacterial susceptibility to cell wall-targeting antibiotics, such as β-lactams. Both depletion and unregulated accumulation of c-di-AMP significantly reduce bacterial resistance to β-lactams. However, despite extensive research on various bacteria (5-11), the molecular mechanisms by which c-di-AMP influences bacterial β-lactam susceptibility remain elusive. Our study is specifically focused on the effects of c-di-AMP accumulation on *L. monocytogenes* β-lactam susceptibility. We identify a previously unrecognized link between c-di-AMP homeostasis, MreB activity, and cell division in *L. monocytogenes*. Through a forward genetic screen, we identified that mutations in MreB, the bacterial rod-shaped determinant protein (12, 14-16), consistently suppress the β-lactam sensitivity of the Δ*PDE* mutant, which accumulates high levels of c-di-AMP. These findings provide mechanistic insight into the cell wall defects observed upon c-di-AMP accumulation in *L. monocytogenes*, and suggest that dysregulation of cytoskeletal dynamics, associated with c-di-AMP accumulation, contributes to impaired cell wall homeostasis and virulence attenuation.

A key observation of this study is the repeated isolation of MreB mutations among suppressor mutants that restore Δ*PDE* resistance to β-lactam antibiotics while still retaining high intracellular c-di-AMP levels. This strongly supports a model in which c-di-AMP accumulation perturbs cell wall physiology, either downstream or independently of its canonical roles in regulating osmotic turgor pressure. Unlike other Firmicutes, which have linked high c-di-AMP levels to increased β-lactam resistance by reducing turgor pressure, our findings strongly suggest that *L. monocytogenes* responds differently, showing increased sensitivity. Interestingly, our study also indicates that the effect of MreB suppressor mutations is independent of potassium (K^+^) homeostasis, a major pathway regulated by c-di-AMP in many bacteria (11). This further suggests that the toxic effects of c-di-AMP accumulation on the cell wall, in *L. monocytogenes*, are largely decoupled from its role in osmotic regulation. Instead, our data point to a model in which c-di-AMP more directly affects cell wall homeostasis, possibly by regulating PG precursor synthesis or enzyme activity. Importantly, the ability of MreB mutations or pharmacological inhibition by A22 to rescue β-lactam sensitivity indicates that MreB activity is detrimental under high c-di-AMP conditions.

Based on the location and nature of the MreB mutations identified, we hypothesize that these mutations likely reduce MreB activity by impairing its polymerization ability and/or its ability to stably associate with the cell membrane. These mutations are located in functionally critical regions of MreB, such as the ATP-binding pocket (G108E), the protomer–protomer interaction interface (A46V), and the RodZ interaction surface (V47I) as indicated by structural mapping (29, 30). These MreB regions are required for ATP-dependent filament assembly, filament stability, and the coupling of MreB activity to the Rod complex (29, 30). Therefore, mutations at these key sites are likely to weaken ATP binding/hydrolysis, disrupt MreB filament formation, or reduce MreB interactions with components of the rod complex, such as RodZ, thereby resulting in less stable or less processive MreB filaments and, consequently, reduced overall MreB activity. Consistent with this, suppressor mutants S3, S6, and S8 exhibit widened cell morphology—a hallmark of reduced MreB function (31)—and phenocopy sublethal treatment with the MreB inhibitor A22 (29). Based on these observations, we hypothesize that the identified mutations act as partial loss-of-function alleles, reducing or dampening MreB activity rather than completely abolishing it. This reduced MreB activity then rebalances cell wall synthesis under conditions of elevated c-di-AMP and reduces the β-lactam sensitivity of the Δ*PDE* mutant.

However, another important implication of our findings is the context-dependent nature of MreB function. While reduced MreB activity is beneficial under high c-di-AMP conditions, our study demonstrates that it is detrimental at normal c-di-AMP levels. Under these conditions, suppressor mutants exhibit abnormal morphology and also do not gain β-lactam resistance. This highlights the need for a delicate balance in cytoskeletal function, where both insufficient and excessive MreB activity can impair bacterial cell fitness, underscoring the importance of regulatory networks that coordinate cytoskeletal dynamics with metabolic and environmental cues.

Our data further suggests that c-di-AMP accumulation disrupts the balance between cell division and elongation. Under β-lactam stress, the Δ*PDE* mutant exhibits pronounced defects in septum formation, leading to excessively elongated, division-impaired cells. This phenotype resembles that of cells deficient in division-specific penicillin-binding proteins, particularly PBP B2, as shown in prior studies (38). Consistent with this, overexpression of PBP B2 (Lmo2039) restores both β-lactam resistance and septum formation/division in the Δ*PDE* background. Together, these observations support a model in which elevated c-di-AMP preferentially impairs the cell division machinery, potentially by limiting the availability or activity of division-specific PG synthesis enzymes. MreB appears to play a crucial modulatory role within this framework. MreB, through the Rod complex, spatially organizes lateral peptidoglycan synthesis, and this activity must be tightly coordinated with cell division (14-16). As previously mentioned, cell widening observed in suppressor mutants, along with the β-lactam resistance conferred by A22 treatment, indicates that reduced MreB activity alleviates the detrimental effects of elevated c-di-AMP (31). One possible explanation is that at high c-di-AMP levels, MreB-driven elongation becomes unbalanced, exacerbating competition for limited PG precursors and thereby depriving the division machinery of them. By dampening MreB activity, suppressor mutations may rebalance PG precursor allocation, allowing sufficient resources for septum formation under stress conditions.

Supporting this model, our metabolic labeling experiments reveal that Δ*PDE* cells exhibit reduced incorporation of both ^14^C-GlcNAc and EDA-DA, i.e., reduced transglycosylation and transpeptidation, respectively. This indicates reduced PG precursor availability in the Δ*PDE* mutant, further depriving the division machinery of them. These experiments also reveal that MreB mutations partially restore both transglycosylation and transpeptidation activities specifically during β-lactam stress. Notably, autolysin activity remains elevated in Δ*PDE* and is not corrected by MreB mutations, suggesting that the rescue of cell wall integrity is not due to reduced hydrolysis but rather improved synthesis. These findings further reinforce the idea that MreB mutations restore a functional balance in peptidoglycan metabolism.

In conclusion, we propose a model in which c-di-AMP accumulation disrupts cell division by impairing septal peptidoglycan synthesis, while continued MreB-driven elongation exacerbates this imbalance. Mutations that reduce MreB activity restore coordination between elongation and division, thereby rescuing cell wall integrity and antibiotic resistance. This work highlights the interplay between second messenger signaling, cytoskeletal dynamics, and cell wall biogenesis, and suggests that targeting this balance may represent a novel strategy for antimicrobial intervention.

## MATERIALS AND METHODS

### Bacterial strains and culture conditions

*L. monocytogenes* and *Escherichia coli (E. coli)* strains in this study are listed in **Table S1**. All *L. monocytogenes* strains were grown in Brain Heart Infusion (BHI) broth or *Listeria* synthetic media (LSM), and *E. coli* strains in Lysogeny broth (LB), unless otherwise indicated, with appropriate antibiotics at 37°C and agitation. For solid media, 1.5% weight/volume agar was added.

### Antibiotic susceptibility assays in BHI

Antibiotic susceptibility in BHI was assessed by measuring bacterial growth in 96-well plates containing 200 µL BHI alone or BHI with two-fold dilutions of cefuroxime, D-cycloserine, moenomycin, bacitracin, and lysozyme. For experiments in which *L. monocytogenes* cultures were treated with MreB inhibitor A22, BHI broth was supplemented with 50 µg/mL A22 (Cayman) prior to the addition of two-fold dilutions of cefuroxime. 2 µL of overnight cultures grown in BHI with shaking at 37°C were inoculated into each well, and bacterial growth was measured by OD_600_ at 37°C with intermittent shaking for 14 hours in a plate reader. For each strain, growth rates at each antibiotic concentration were normalized to the growth rate in BHI only.

### Antibiotic susceptibility assays in LSM Broth

Antibiotic susceptibility in LSM was assessed by growing bacterial cultures in sodium-buffered LSM (Na-buffered LSM) supplemented with either 0.1 mM, 4.8 mM, or 10 mM KCl, as indicated. For all experiments, starter cultures were grown in 5 mL Na-buffered LSM containing 4.8 mM KCl in 125-mL baffled flasks at 37 °C with shaking to mid-log phase (OD600 ∼ 0.5). Bacterial cells were then harvested and washed once with Na-buffered LSM containing 0.1 mM KCl, then resuspended in 1 mL of Na-buffered LSM supplemented with 0.1 mM KCl. OD_600_ was measured, and cultures were normalized to an OD_600_ ∼ 0.5. From these normalized cultures, 20 µL was used to inoculate 1 mL of fresh Na-buffered LSM containing either 0.1 mM, 4.8 mM, or 10 mM KCl, in the presence or absence of cefuroxime (3 µg/mL). Cultures were incubated at 37 °C with shaking, and growth was assessed by measuring OD_600_ after 16 hours. For each strain and each KCl concentration, growth at each antibiotic concentration was normalized to the growth of the same strain in LSM only.

### Spot dilution growth assay on Na-Buffered LSM agar

Overnight cultures of bacterial strains were initiated in Na-buffered LSM supplemented with 4.8 mM KCl and grown at 37 °C with shaking. The following day, overnight cultures were diluted into fresh Na-buffered LSM medium with 4.8 mM KCl to initiate daytime cultures. These cultures were grown at 37 °C with shaking until mid-log phase, i.e., an OD_600_ ∼ of 0.5. Cultures were then normalized to an OD_600_ ∼ 1.0 using Na-buffered LSM containing 0.1 mM KCl. Ten-fold serial dilutions were prepared in the same medium, and 5 µL of each dilution was spotted onto LSM agar plates supplemented with or without 3 µg/mL of cefuroxime. For untreated conditions, dilution series ranged from 10^-3^ to 10^-8^, whereas for cefuroxime-containing plates, dilution series ranged from 10^0^ to 10^-5^. Plates were incubated at 37 °C, and growth was assessed following incubation.

### Mammalian L2 plaque assay

L2 fibroblast cells were grown in DMEM high glucose (Sigma) with 1% of sodium pyruvate and 1% of L-glutamine in the presence of 10% fetal bovine serum (FBS). Plaque formation upon *L. monocytogenes* infection of L2 cells was quantified as previously described. Briefly, 1.2 × 10^6^ L2 cells were infected with *L. monocytogenes* at a multiplicity of infection (MOI) of 0.25. At 1-hour post-infection, cells were washed with 1X TC-grade PBS, and fresh cell medium containing 0.7% agarose was supplemented with 10 μg/mL gentamicin to kill extracellular *L. monocytogenes*. At 5 days post infection, cells were stained with 0.3% crystal violet to visualize plaques. Plaque sizes (in mm^2^) were analyzed using ImageJ software. For each strain, the plaque sizes of each plaque were normalized to the average plaque size of wild-type *L. monocytogenes*.

### Fluorescence Microscopy

*L. monocytogenes* strains were grown in either plain BHI broth or BHI with 0.5X MIC of cefuroxime to mid-log phase, i.e., an OD_600_ ∼ 0.5. Bacterial cells were harvested by centrifugation, washed twice with 1X PBS, and resuspended in 1X PBS. Next, 100 μL of cell suspension was incubated with 10 μg/mL WGA for 10 min in the dark at room temperature. After incubation, 1.5μL of cell suspension was pipetted onto an agarose pad (1% agarose in PBS, placed on a microscopy slide), and sealed under a coverslip. Bacterial cells were imaged with a Nikon A1R Confocal microscope (Nikon, Tokyo, Japan) at 100x magnification with a gain of 60. The bacterial cell length and width were measured using the MorphoSegger program, created by Andres Florez on GitHub. Between 300 and 500 cells were quantified per condition across three independent experiments. Quantitative length and width measurements were compiled into a single dataset and analyzed by frequency distribution. Values were grouped into fixed-length bins of 0.5 μm, spanning 0-10 μm. Similarly, values were grouped into fixed-width bins of 0.25 μm, spanning 0-2.25 μm. For cells treated with A22, cell width values were grouped into 0.05-μm-wide bins. Each data point was assigned to exactly one bin, and exact counts were calculated for each bin, which were used for downstream analysis and visualization.

### ^14^C-GlcNAc incorporation

*L. monocytogenes* strains were grown in BHI or BHI with 0.5x MIC of cefuroxime to mid-log phase, i.e., an OD_600_ ∼ 0.5. 0.32μL of ^14^C-GlcNAc was added to 120μL of culture for a final concentration of 0.27mCi/mL. The sample was then incubated with shaking at 37°C. At time points 15min and 30min, the cells were collected on a 0.25 μm filter paper and washed with 1X PBS. The filter paper, as well as bacterial cells, were transferred into a pony vial containing 3 mL of scintillation fluid, and the prepared sample will be analyzed for ^14^C-GlcNAc using a liquid scintillation analyzer (Tri-Carb 4910TR) as previously described. Radioactivity uptake was measured as counts per minute (CPM), and the raw radioactivity values (CPM) were normalized to the corresponding OD_600_ values measured at the same time points to obtain absolute ^14^C-GlcNAc uptake.

### EDA-DA incorporation

*L. monocytogenes* strains were grown in BHI or BHI with 0.5x MIC of cefuroxime to mid-log phase, i.e., an OD_600_ ∼ 0.5. At this point, culture was transferred to a fresh culture tube, and EDA-DA was added to a final concentration of 0.5 mM. Cultures were incubated at 37 °C with shaking, and samples were collected at 15 min and 30 min. Cells were washed with 1X PBS, fixed in pre-chilled 70% ethanol at −20 °C for 10 min, and washed with 1X PBS containing 3% BSA. Fixed cells were subjected to click-chemistry labeling using Alexa Fluor-PCA, incubated for 30 min at room temperature in the dark, and washed again with 1X PBS containing 3% BSA. Cells were resuspended in 1X PBS, and OD_600_ was measured using the plate reader for normalization. Raw fluorescence was measured from appropriately diluted cell suspensions in a 96-well plate using the plate reader, using excitation at 488 nm, emission at 520 nm, and a gain of 100, and values were normalized to OD600 to obtain absolute fluorescence.

### Triton X-100–Induced Lysis

*L. monocytogenes* strains were grown to mid-log phase (OD_600_ ∼ 0.5) in BHI or BHI + 0.5x MIC of cefuroxime. The bacterial cells were then washed with 1X PBS and resuspended in 0.1% volume/volume Triton X-100 in phosphate-buffered saline to a final OD_600_ ∼ 1. 200 μL of bacterial suspension was transferred into a 96-well plate, and the cell lysis was monitored by OD_600_ measurements in a plate reader.

