## Supplementary material for "Mutations in MreB suppress β-lactam sensitivity upon c-di-AMP accumulation in *Listeria monocytogenes*": Figures S1-S4

**Table S1:** Strains used in this study

| Strain number | Genotype | Source |
|---|---|---|
| ***L. monocytogenes*** (derivatives of strain 10403S) | | |
| TNHL261 | WT | (1) |
| TNHL22 | Δ*pdeA* Δ*pgpH (*Δ*PDE)* | (2) |
| TNHL376 | S1 | This study |
| TNHL392 | S2 | This study |
| TNHL393 | S3 | This study |
| TNHL394 | S4 | This study |
| TNHL418 | S5 | This study |
| TNHL414 | S6 | This study |
| TNHL416 | S7 | This study |
| TNHL417 | S8 | This study |
| TNHL395 | S9 | This study |
| TNHL396 | S10 | This study |
| TNHL29 | Δ*PDE*+ $P_{spac}$-*pgpH* | (3) |
| TNHL526 | S1 + P*spac-pgpH* | This study |
| TNHL527 | S2 + P*spac-pgpH* | This study |
| TNHL528 | S3 + P*spac-pgpH* | This study |
| TNHL529 | S4 + P*spac-pgpH* | This study |
| TNHL536 | S5 + P*spac-pgpH* | This study |
| TNHL532 | S6 + P*spac-pgpH* | This study |
| TNHL534 | S7 + P*spac-pgpH* | This study |
| TNHL535 | S8 + P*spac-pgpH* | This study |
| TNHL530 | S9 + P*spac-pgpH* | This study |
| TNHL531 | S10 + P*spac-pgpH* | This study |
| TNHL1166 | WT + P*spac-lmo2039* | This study |
| TNHL1167 | Δ*PDE* + P*spac-lmo2039* | This study |
| ***E. coli*** | | |
| TNH47 | pPL2-Pspac-*pgpH* in *XL1B* | This study |
| TNH1220 | pPL2-Pspac-hly-5'UTR-*lmo2039* in *DH5α* | This study |

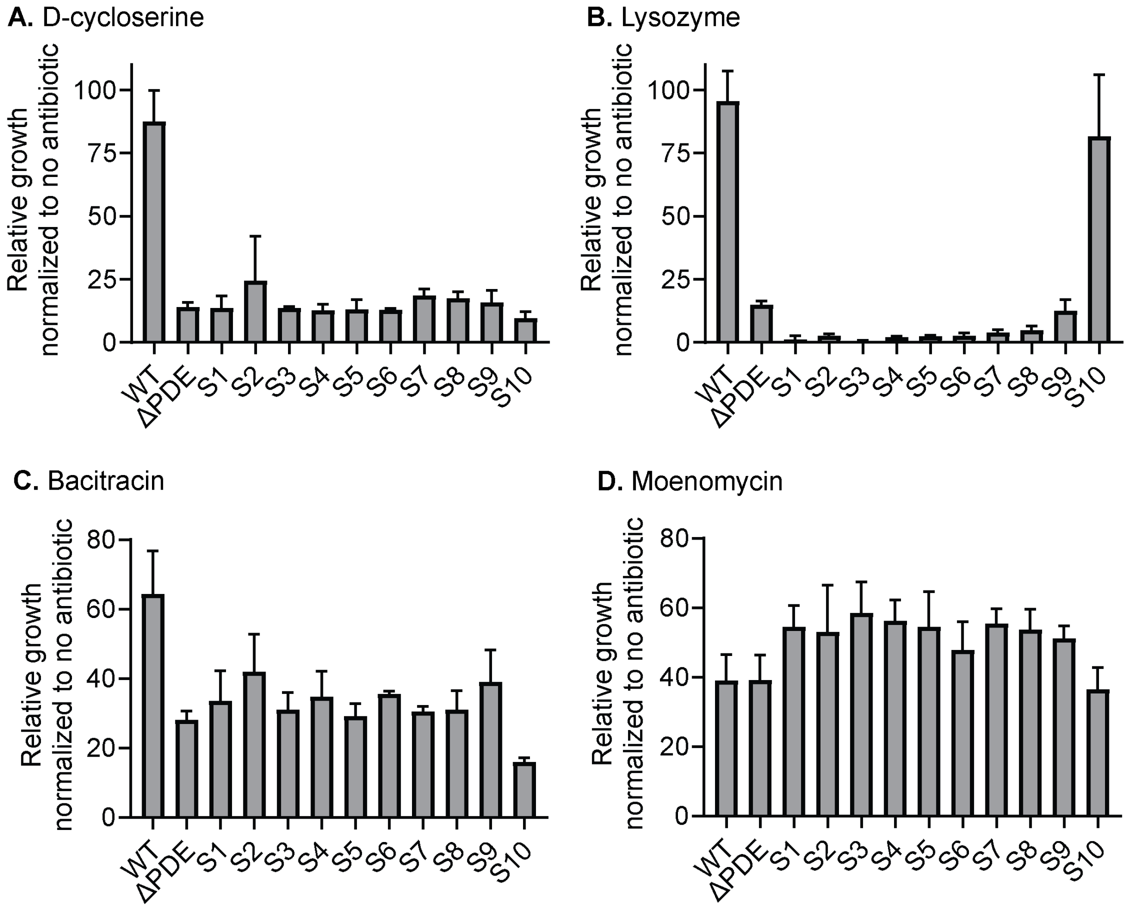


**Figure S1: Sensitivity of *L. monocytogenes ΔPDE* and suppressor mutants to cell wall-targeting antimicrobials other than β-lactams.** Relative growth rates of *L. monocytogenes* in BHI with 12.5 μg/mL D-cycloserine **(A)**, 0.5 mg/mL lysozyme **(B)**, 125 μg/mL bacitracin **(C)**, and 30 ng/mL moenomycin **(D)**, normalized to growth rates in BHI only for each strain.

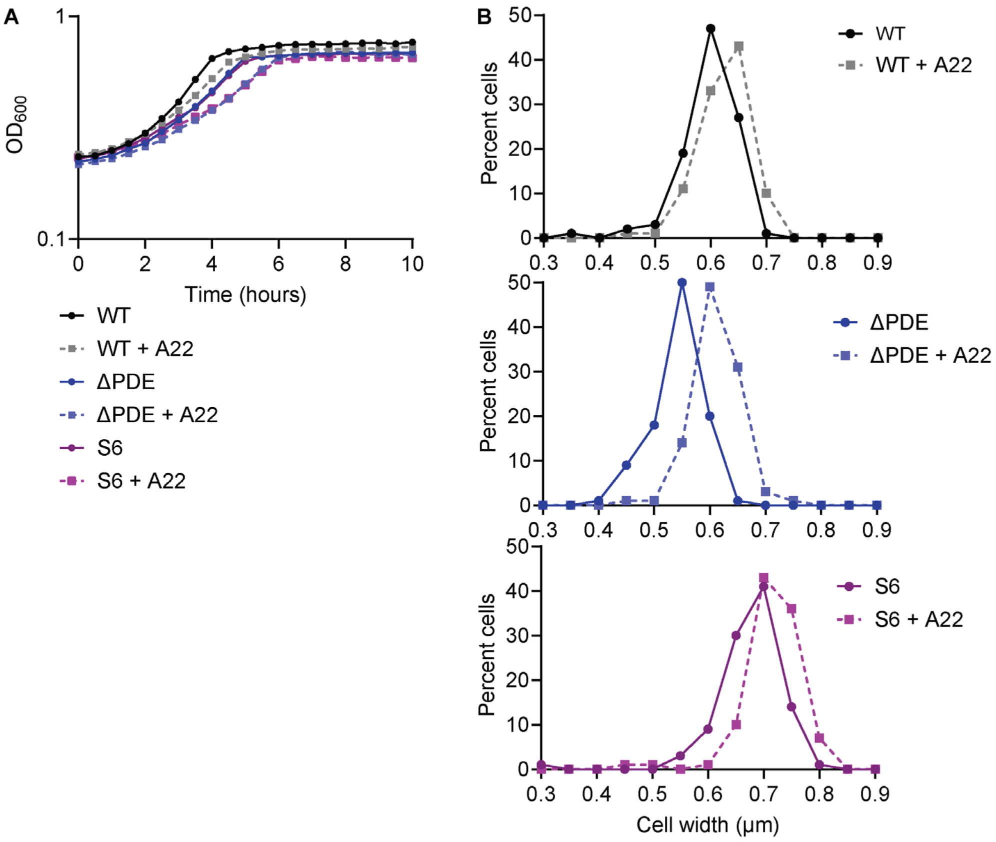


**Figure S2: A22 treatment widens *L. monocytogenes* WT and Δ*PDE* mutant. A.** Growth of *L. monocytogenes* in BHI or BHI + 50 µg/mL A22. **B.** Cell widths distribution upon A22 treatment. Around 100 cells were quantified per condition across three independent experiments. Width measurements were compiled into a single dataset and analyzed using a frequency distribution, with values grouped into 0.05-µm-wide bins. Bin boundaries were defined as left-inclusive and right-exclusive, and each data point was assigned to exactly one bin. Exact counts were calculated for each bin.

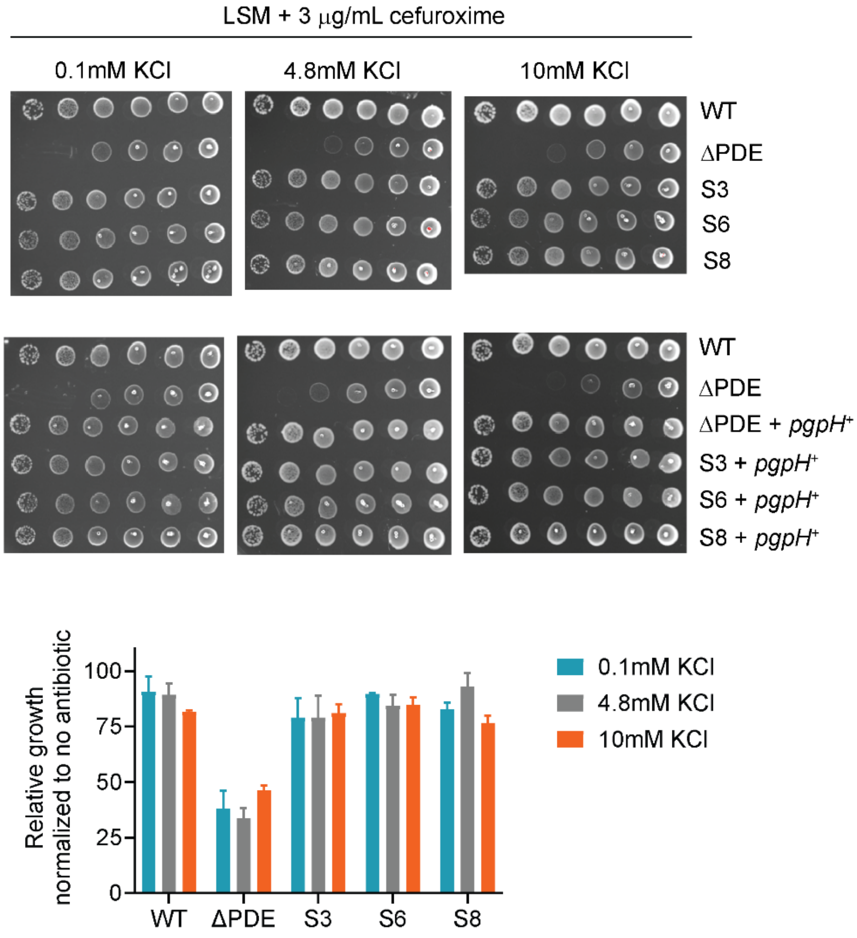


**Figure S3: The *L. monocytogenes* ΔPDE mutant is β-lactam sensitive, independent of $K^+$ availability. A. Spot dilution on LSM agar containing different $K^+$ concentrations. Top:** Mid-log cultures were normalized to the same density and spotted onto LSM agar with 3 μg/mL of cefuroxime at varying $K^+$ concentrations. **Bottom:** Cultures were grown in LSM with varying KCl concentrations, and with or without 3 μg/mL cefuroxime. Growth in cefuroxime was normalized to growth in LSM only containing the same $K^+$ concentration. Data are the average of 5-6 independent experiments. Error bars represent standard deviation.

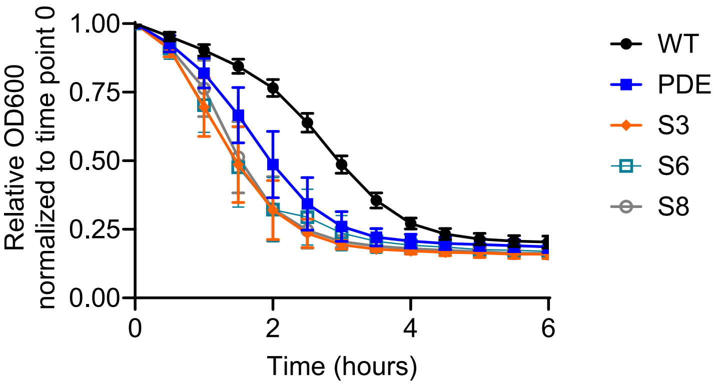


**Figure S4: ΔPDE has increased autolysin activity, while MreB mutations have a modest effect on it.** *L. monocytogenes* strains were grown to mid-log phase ($OD_{600}$ ~0.5) in BHI, and lysis was induced by 0.1% Triton X-100.
